# An intermittent hypercaloric diet alters gut microbiota, prefrontal cortical gene expression and social behaviours in rats

**DOI:** 10.1101/328294

**Authors:** Amy C Reichelt, Amy Loughman, Ashton Bernard, Mukesh Raipuria, Kirsten N Abott, James Dachtler, Thi Thu Hao Van, Robert J Moore

## Abstract

Excessive consumption of high fat and high sugar (HFHS) diets are known to alter reward processing and aspects of behaviour, and change microbiota profiles. Studies in gnotobiotic mice also provide evidence that gut microorganisms influence social behaviour. To further investigate these interactions, the impact of intermittent access to a HFHS diet on social behaviour, gene expression and microbiota composition was examined. Rats were permitted intermittent daily access (2h / day) to a palatable HFHS diet for 28 days across the adolescent period. Social interaction, social memory and novel object recognition were assessed during this period. Following testing, RT-PCR was conducted on hippocampal and prefrontal cortex (PFC) samples. 16S ribosomal RNA amplicon sequencing was used for identification and relative quantification of bacterial taxa. Reduced social interaction behaviours, and impaired social memory and novel object recognition were observed in HFHS diet rats. Reduced levels of monoamine oxidase A (Maoa), catechol-O-methyltransferase (Comt) and brain derived neurotrophic factor (Bdnf) mRNA were observed in the PFC of HFHS diet rats. The relative abundance of a number of specific taxa differed significantly between the two diet groups, in particular, *Lachnospiraceae* and *Ruminoccoceae* bacteria, which also predicted social behaviours, novel object recognition performance and Maoa expression. This is the first study to show that limited daily access to HFHS diet alters social behaviour and cognition in rats. Furthermore, behavioural changes are associated with alterations to cortical gene expression of enzymes involved in monoamine synthesis and neuroplasticity, and microbiota profiles predicted diet-induced changes to behaviour and gene expression.

## 1. Introduction

The global rate of obesity is rapidly growing, and the incidence of overweight and obesity is particularly increasing amongst young people and children (Ogden et al., 2014). Epidemiological studies indicate that adolescents and young adults most frequently consume hypercaloric high fat and high sucrose (HFHS) “junk” foods (Braithwaite et al., 2014), increasing negative health outcome risks.

Chronic exposure to hypercaloric diets causes multiple changes to behavioural processes and neuromodulation within the brain, which has been linked to decreased dopamine turnover in the mesolimbic system (Davis et al., 2008). However, the effects of chronic HFHS diet consumption may mask critical periods in development whereby intermittent consumption of this diet leads to lasting biological and behavioural changes in adulthood. Indeed, emerging data suggests that adolescence may be a sensitive period for susceptibility to diet-induced behavioural changes in mood (Baker et al., 2017), reward seeking (Naneix et al., 2017; Reichelt, 2016) and cognition (Labouesse et al., 2017).

Beyond a role in cognition, recent studies have suggested links to obesity, hypercaloric diet consumption and changes in social behaviour in rodents (Carvalho et al., 2016; Teixeira et al., 2017; Yaseen et al., 2018). High fat diet consumption increased social interaction in adult male mice, but impaired recognition memory of a novel mouse (Takase et al., 2016), and social recognition has also been recently shown to be altered in juvenile rats following short term exposure to high fat diets (Yaseen et al., 2018). Social play is a characteristic adolescent social behaviour that decreases into adulthood (Trezza et al., 2010). Social play was reduced following neonatal overfeeding, suggesting that nutrition may impact the expression of this behaviour (Carvalho et al., 2016); however litter size manipulations may contribute to altered social repertoires.

There are links between the brain regions that support social behaviour and those that are altered by HFHS diet. The prefrontal cortex (PFC) matures across adolescence (Spear, 2000) and represents a critical period of vulnerability to diet evoked cognitive deficits (Baker et al., 2017; Reichelt and Rank, 2017). The PFC has an important role in social processing (Bicks et al., 2015; Kolb, 1974), and appropriate maturation is fundamental for the development of social cognition (Kim et al., 2015). Experimental evidence highlights that the rodent homologue of the medial PFC and the hippocampus are important for social behaviour, including social memory and sociability (Kogan et al., 2000; Okuyama et al., 2016; Rudebeck et al., 2007). Aspects of social interaction are rewarding (Trezza et al., 2010), and the increased dopamine neurotransmission and refinement of reward-associated neural connections within the PFC across adolescence is proposed to invigorate this behaviour.

Moreover, previous observations noted that intermittent access to a HFHS diet (Baker and Reichelt, 2016), or high fat diet (Labouesse et al., 2017) across adolescence evoked PFC dysregulation, complementing research demonstrating that PFC excitation/inhibitory imbalance underpins social deficits (Selimbeyoglu et al., 2017), thus providing rationale for exploring the impact of an intermittent HFHS diet on social behaviours. Restricted access to palatable foods has been shown to impact on reward neurocircuitry (Bocarsly et al., 2014; Furlong et al., 2014), and furthermore allows behavioural examination both immediately following palatable food consumption and when animals have not had access to palatable foods.

Diet influences the gut microbial composition (Albenberg and Wu, 2014) and alterations to gut flora has been linked to changes in cognition, mood and behaviour (Desbonnet et al., 2015; Frohlich et al., 2016). Studies utilising germ-free (GF) mice demonstrate that the presence, composition, and functionality of the gut microbiota is crucial for normal social behaviours, which are reduced in GF mice (Desbonnet et al., 2015). Moreover, GF mice and antibiotic-induced gut dysbiosis rodent models have demonstrated associations between disruption of the gut microbial community and cognitive, social and emotional alterations (Desbonnet et al., 2015; Frohlich et al., 2016).

To further the evidence that intermittent exposure to a HFHS diet during the juvenile developmental phase alters cognitive control and neurotransmitter systems within the brain, we examined the effects of intermittent HFHS food consumption on social interaction and social memory in young rats. We highlight putative molecular pathways by examination of the expression of genes associated with neuroplasticity, monoamine signalling, and neuroinflammation in the PFC and hippocampus, and faecal microbiota composition to explore diet-induced alterations. Spontaneous novel object recognition and odour recognition memory were examined to assess HFHS diet effects on memory and olfaction. Exploratory analyses through linear modelling were performed to determine associations between faecal microbiota composition, behaviour and cortical gene expression.

## 2. Methods

### 2.1. Animals

Male (*n*=32) albino Sprague Dawley rats (Animal Resources Centre, Western Australia) arrived at postnatal day (P)21 (~50g) and were housed in groups of four in climate (21°C ± 2°C; humidity 55 ± 5%) and light (12 h cycle lights on at 07:00h) controlled colony room. Standard laboratory rat chow (Meat Free Rat and Mouse Diet, Specialty Feeds, Western Australia; energy composition of 14 KJ/g, 23% protein, 12% fat, 65% carbohydrates) and water was available *ad libitum* throughout the experiment. Behavioural tests were performed between 08:00 and 14:00h and procedures were approved by the Animal Care and Ethics Committee at RMIT University.

### 2.2. Diet administration

Rats were allocated to diet conditions: Control (chow fed, *n*=8) or HFHS condition (*n*=8), or were allocated as age/weight matched sample for social memory and social interaction (*n*=16). Body weights were standardized in all treatment groups prior to the commencement of the diet (Control: 75.5 ± 2.0g; HFHS: 76.4 ± 2.0 g), and rats were handled for 7 days prior to manipulations. Group-housing was used to negate social isolation stress (Skelly et al., 2015). Rats in the HFHS diet condition were provided with 2 h daily homecage access (between 09:00-11:00h) to semi-pure HFHS pellets (Specialty Feeds, Western Australia, SP04-025; 18.4kJ/g digestible energy; composed of 20% fat (lard), 39.6% sucrose, 19.4% protein, providing 36% energy from lipids and 55% from sucrose), in addition to *ad libitum* chow (Specialty Feeds, Western Australia, Meat Free Rat and Mouse Diet, energy composition of 14 KJ/g, 23% protein, 12% fat, 65% carbohydrates) and water access. Consumption of HFHS diet was calculated from the weight (g) between the HFHS pellets allocated and that collected after 2 h access per cage. Body weight was recorded at baseline before the diet began, and thereafter twice per week. Chow consumption over a 24 h period was measured twice per week in conjunction with HFHS pellet intake to calculate total energy intake per cage of four rats (Del Rio et al., 2016).

### 2.3. Behavioural analysis

A timeline of the general experimental procedures is illustrated in Figure 1A. Diet administration began on P28, coinciding with definitions of adolescence in male rats (Spear, 2000). Behavioural tests were conducted in a room illuminated to 30 lux, rats were assessed for social interaction, social memory, social odour preference, novel object recognition and odour recognition memory. All behavioural data were scored by an observer who was blind to the group allocations using ODLog (version 2.7, Macropod Software, Australia).

#### 2.3.1. Social interaction

Social interaction tests were conducted in a square test arena (dimensions: 50 cm [length] x 50 cm [width] x 60 cm [height]) constructed from black Perspex. A camera mounted above the test area recorded all the social interaction tests to a computer for subsequent scoring. All rats were habituated to the arena 24 h prior to testing by being placed individually into the arena for 10 minutes.

Prior to social interaction testing, rats were isolated from their cage mates in individual holding cages for 15 minutes. In the social interaction test, one rat from either the control or HFHS diet condition rat was placed in the arena with an unfamiliar partner matched for body weight (+/− 10g). Test sessions were 10 min duration. To differentiate between animals, one rat was marked on its back with a black odourless fabric pen marker 24 h prior to testing. The two rats were placed into the test arena simultaneously so that they were facing each other in opposing corners. Rats in the HFHS diet condition were tested 1 h after access to the HFHS pellets “post”, and 23 h after HFHS pellet access “pre”, counterbalanced across days and animals. Between tests the arena was cleaned with 70% ethanol to eliminate odour cues.

As social behaviour in rats has been shown to depend on the playfulness of its partner, both animals in a sample pair were considered as one experimental unit (Trezza et al., 2010). Pinning and pouncing frequencies were quantified and considered the most characteristic parameters of social play behaviour in rats. Social play behaviours usually occur very rapidly and they are of short duration thus, individual frequency was scored. Videos were scored to measure i) the total time (sec) spent in social interaction, ii) frequency of social investigation behaviour (sniffing, licking, grooming), iii) frequency of social play behaviour (pinning, pouncing), iv) frequency of aggressive-like behaviour (rump biting, boxing, overt physical harm).

#### 2.3.2. Social memory

Social memory was tested in two phases (see Fig 1D). As HFHS rats showed differences in social interaction pre HFHS food consumption, social memory testing and other behavioural tests were conducted after HFHS access to ensure that any memory deficits observed were not due to reduced social contact in the HFHS diet rats. Social memory tests were conducted in a circular arena (dimensions: 100 cm diameter, 50 cm high) constructed from grey Perspex. The arena contained two wire chambers with plastic bases (dimensions: 18 cm [length] × 20 cm [width] × 22 cm [height]). The wires were interspaced 1cm apart which allows the test rats to interact with the novel sample rats but not physically contact them. A camera mounted above the test area recorded the tests as described above. Control and HFHS diet rats were habituated to the testing apparatus 24 h prior to testing by being placed individually into the arena with the empty chambers for 10 minutes. Sample rats were also habituated to the individual chambers for 10 minutes 24 h prior to testing.

Social memory was tested in two phases. In Phase 1, rats were placed in the arena for 5 min with one sample rat in a chamber and the other chamber empty. Time exploring the chamber containing the sample rat versus the empty chamber was considered a measure of sociability (Crawley et al., 2007). The experimental rat was then removed and placed into individual holding cages for a 5 min inter-trial interval (ITI) period. In Phase 2, the arena contained the original sample rat (familiar) in a chamber and the previously empty chamber contained a novel rat. The experimental rat was returned to the arena to explore for a 3 min period. Between test phases the arena was cleaned with 70% ethanol to eliminate odour cues. Videos were scored to measure the duration of time the rat spent exploring the chambers during each phase. Sociability was quantified as the time spent exploring the chamber containing the sample rat as opposed to the empty chamber, and social recognition memory was measured as the time spent in proximity to the chamber containing the novel rat versus the familiar sample rat.

#### 2.3.3. Social odour preference

The chambers used for social recognition were filled with soiled bedding from a cage of young male rats (approximately 5 weeks of age) housed in an adjacent holding room, or clean bedding. Rats were allowed to freely explore the arena for 5 min and the amount of time spent exploring empty chambers containing either soiled or clean bedding was recorded.

#### 2.3.4. Odour memory

Odour memory was conducted in the square test arena (as described in 2.3.1). Identical cylindrical stainless steel containers (10cm [height] × 6cm [width]) with perforated stainless steel lids were filled with corncob bedding and then scented with 3 ml of peppermint or almond extract (Queen, Australia) to serve as odour stimuli (see Fig 1E). The odour memory test consisted of 2 phases – sample and test. Pilot testing determined that these odours were equally explored by the rats. During the sample phase two of the same scented containers were placed in opposite corners of the arena. The rat was allowed to freely explore the arena for 5 min. The rat was then removed from the arena and placed in a holding cage for a 5 min retention period. The arena was thoroughly cleaned with 70% ethanol and one of the scented containers was replaced with an identical container filled with a novel odour. The rat was then returned to the arena for a 3 min test phase. The duration of time the rat spent exploring each of the odour containers during each phase was measured.

#### 2.3.5. Object recognition memory

Object recognition (Fig 1F) was conducted in the square test arena (as described in 2.3.1). Commercial objects (e.g. plastic bottles and tin cans) were used with differing heights (16-24cm) and widths (7-14cm). Rats explored two identical sample objects in the arena (sample phase; 5 minutes). The following day, 24 h after the sample phase, rats were tested for recognition of a familiar versus a novel object (test phase; 3 mins). The duration of time the rat spent exploring each object during each phase was measured.

### 2.4. Sample collection

Rats were anaesthetised (sodium pentobarbital 100 mg/kg, intraperitoneal), brains removed and the PFC and hippocampus (composed of dorsal and ventral poles) dissected and snap frozen in liquid nitrogen and stored at −80°C for analysis by RT-PCR. Retroperitoneal and gonadal white adipose tissues (rpWAT; gnWAT) and liver were dissected and weighed. Livers were visually scored for markers of hepatic steatosis based on previous criteria (Velkoska et al., 2010). One faecal bolus was collected from the terminal caecum, snap frozen and stored at −80°C prior to microbiota analysis.

### 2.5. Quantitative RT-PCR

RNA was extracted using Tri-Reagent (Sigma-Aldrich) and RNeasy Mini kit (Qiagen), and quantity and purity of RNA determined by nanodrop. RNA was converted to cDNA (RT^2^ First Strand Kit Qiagen). Gene expression was quantified by Custom RT^2^ Profiler PCR Arrays (Qiagen) with RT^2^ SYBR Green Mastermix (Qiagen, Australia), real-time PCR was then performed using a QuantStudioTM 7 Flex Real-Time PCR System (Applied Biosystems). Genes of interest were NLR family, pyrin domain containing 3 (Nlrp3), Glutamate decarboxylase 1 (Gad1), Brain-derived neurotrophic factor (Bdnf), Dopamine receptor D1 (Drd1), Dopamine receptor D2 (Drd2), Monoamine oxidase A (Maoa), Catechol-O-methyltransferase (Comt), 5-hydroxytryptamine (serotonin) receptor 4, G-coupled (Htr4), Tumour necrosis factor alpha (Tnf-α), Interleukin 6 (Il6), Integrin, alpha M (Itgam) and Actin, beta (Actb) from Qiagen (See Supplementary Table 1 for reference sequences). Analysis of relative gene expression was normalized to the housekeeping gene beta actin, via the ΔΔC_T_ method (Livak and Schmittgen, 2001).

### 2.6. 16S rRNA gene amplicon sequencing and bioinformatics

Total DNA was isolated using the Bioline ISOLATE Faecal DNA Kit (Bioline). PCR was performed using Q5 DNA polymerase (New England Biolabs) with a primer set selected to amplify V3-V4 region of 16S rRNA gene (forward: ACTCCTACGGGAGGCAGCAG and reverse: GGACTACHVGGGTWTCTAAT). Sequencing was performed on an Illumina MiSeq instrument (2 × 300bp paired-end sequencing), following the method detailed by Fadrosh et al. (2014). Sequences were joined in Quantitative Insights Into Microbial Ecology (QIIME) 1.9.1 (http://qiime.org) using the fastq-join method. Maximum allowed percent differences within the overlapping region was zero. Sequences were de-multiplexed using the QIIME split library protocol, keeping only sequences with Phred quality score higher than 20. The dataset was inspected for chimeric sequences using Pintail (Ashelford et al., 2005). Operational taxonomic units (OTUs) were clustered at 97% sequence identity using UCLUST (Edgar, 2010). Taxonomic assignments were performed against the GreenGenes database (DeSantis et al., 2006). OTUs with a relative abundance of less than 0.01% were removed.

### 2.7. Statistical analyses

#### 2.7.1. Behaviour, physiological parameters and brain mRNA expression

Results were analysed using repeat measures analysis of variance (ANOVA - body weight and energy intake), mixed design ANOVAs (social recognition memory, social interaction, sociability, novel odour recognition and novel object recognition), one-way ANOVA (rpWAT, gnWAT, liver weight, RT-PCR values) with post-hoc Tukey and equality of error variance assessed, or multivariate linear models following significant correlations with post-hoc tests. ΔΔC_T_ values that exceeded ±2 standard deviations from the mean were excluded from analysis, resulting in group sizes of 6-8 per gene.

Social recognition memory and novel object recognition performance were converted to Exploration Ratios [Ratio = Time(novel-familiar) / Time(novel+familiar)] to permit bivariate analysis using correlations (Pearson’s R, 1-tail) to examine associations between mRNA expression and behaviours found to differ between diet groups. Liver scores were analysed using the Kruskal-Wallis test. Data were analysed with IBM SPSS Statistics 24, GraphPad Prism 7 and R.

#### 2.7.2. Microbiota

Visualisation, alpha diversity and distance measures of microbiota were conducted using the R packages *phyloseq* and *MixOmics*. Data were total-sum scaled (i.e. relative abundance of operational taxonomic units – OTUs) and centre-log ratio transformed where appropriate. The *DESeq2* package was used to undertake differential abundance testing (Love et al., 2014), and multivariate analysis of variance (MANOVA) was used to test associations between *Firmicutes* to *Bacteroidetes* (FB) ratio, behaviour and gene expression.

Significance for differential abundance analyses was assessed on the basis of a threshold q-value of 0.05 (i.e. p-value adjusted using the False Discovery Rate approach; Benjamini et al. (2001)). Bivariate correlations were calculated using Pearson’s R, 2-tailed.

## 3. Results

### 3.1. Body Weight, energy consumption and physiological measurements

All rats gained weight across the experiment, however HFHS diet rats gained more weight than controls (Fig 1B x diet group *F*(8,112)=5.07, *P*<0.001). Overall, rats consumed increasing amounts of energy across the 4 week period (*F*(3,18)=81.4, *P*<0.001), and HFHS diet rats consumed more energy than control rats (Fig 1C, diet group x time *F*(1,6)=10.8, *P*<0.001). At the experimental end point, HFHS diet rats were heavier than chow fed animals (*F*(1,14)=4.516, *P*=0.05), had greater rpWAT (*F*(1,14)=5.54, *P*=0.034), gnWAT (*F*(1,14)=4.71, *P*=0.048), and increased liver scores (*U*=5, *P*<0.01) (Supplementary Table 2).

**Figure 1.**
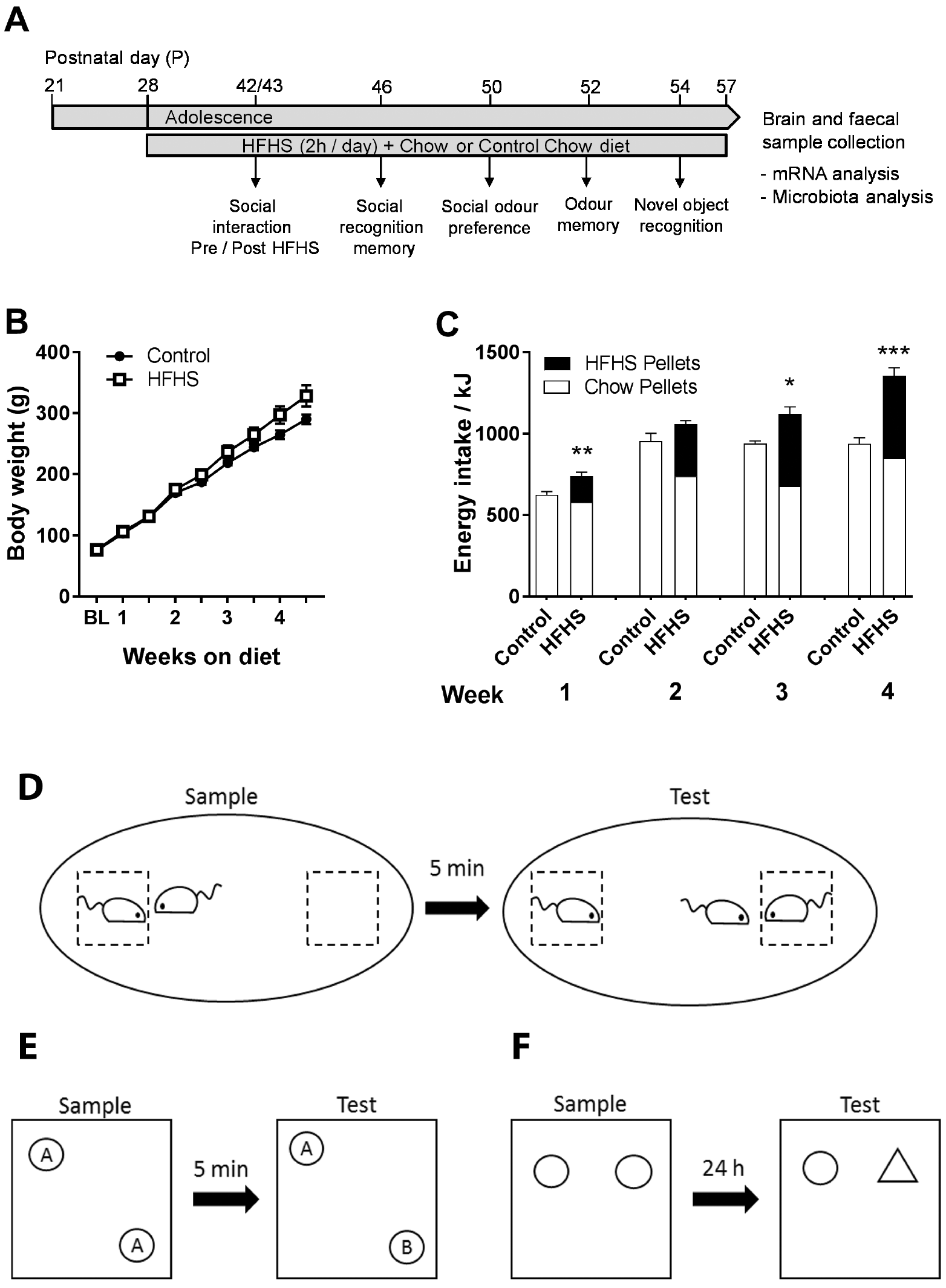
A) Timeline of experimental procedures showing the ages of the rats at each behavioural test (conducted at the same age in all animals) and at sacrifice. Social interaction testing was conducted both before and after access to palatable HFHS food in the HFHS rats. As HFHS rats showed differences in social interaction pre HFHS food, social memory testing, and other behavioural tests were conducted after HFHS access, to ensure that any
memory or behavioural deficits observed were not due to reduced social contact in the HFHS diet rats. B) Mean body weights of control and HFHS rats across the 4-week diet exposure period. C) Mean energy consumption (kJ) per cage of rats across the 4-week diet exposure period. D) Schematic of social memory testing procedure. E) Schematic of novel odour recognition procedure, where A and B are different odours contained in identical containers. F) Schematic of novel object recognition procedure. Error bars represent +SEM. * indicates *P*≤0.05, ** *P*<0.01, *** *P*<0.001.

### 3.2. Effect of HFHS diet on social interaction

To assess the effect of what and when HFHS diet consumption has upon social behaviour, we tested the total social exploration time one hour prior (“pre”) or one hour following (“post”) HFHS food access. Social interaction duration did not differ in the control (normal chow fed) animals. However, HFHS diet rats spent less time engaged in social interaction “pre” HFHS food access, compared to “post” HFHS food access (diet access x diet group *F*(1,14)=5.66, *P*<0.05, effect of diet group “pre” *F*(1,14)=9.271, *P*<0.01, but not “post” F<1, Fig 2A). The microstructure of social behaviour was also examined. Social investigation frequency was increased in the HFHS rats post-consumption (diet access x diet group *F*(1,14) = 8.6, *P*<0.05; HFHS *F*(1,14)=21.59, *P*<0.001, control *F*<1, Fig 2B). No differences were observed in the frequency of social play behaviours (Fig 2C), and aggressive behaviours were not observed. Together, this data suggests that for those rats on the intermittent HFHS diet, social motivation is decreased the longer the period is since diet consumption.

### 3.3. Effect of HFHS diets on social recognition memory

Mouse social behaviour has been typically examined using the ‘three-chamber’ social approach test. We adapted this protocol for use in rats to assay whether changes in social recognition memory could be altered by HFHS diet. During the social approach phase of the sociability test (Fig 2D), both control and HFHS rats preferentially explored the novel rat, “sample”, compared to the empty cage (*F*(1,14)=275.5, *P*<0.001), no significant between group or interaction effect, *F*s<1). However, HFHS rats showed impaired social recognition, exploring the familiar and novel rat equally, contrasting to the strong preference of control rats to explore the novel rat (chamber x diet group *F*(1,14)=39.15, *P*<0.001, control *F*(1,14)=109.3, *P*<0.001, HFHS *F*(1,14)=2.6, *P*=0.13, Fig 2E). Exploration ratios calculated from the test data [Mean (SEM): Control = 0.80 (0.03); HFHS = 0.56 (0.03)] differed significantly between groups (*F*(1,14)=33.2, *P*<0.001).

### 3.4. No effect of diet on social odour preference or odour recognition memory

To confirm that the lack of social recognition memory in the HFHS rats was not due to a lack of olfactory sensitivity, we tested their ability to discriminate between clean and soiled bedding and between two non-social odours. Control and HFHS diet rats preferentially explored the chamber containing a social odour (*F*(1,14)=217.8, *P*<0.001, Fig 2F). During odour recognition testing, control and HFHS diet rats preferentially explored the novel odour container, demonstrating odour recognition memory (odour x diet group *F*(1,14)=3.0, *P*=0.105, Fig 2G). Together, HFHS rats were unimpaired in odour discrimination, suggesting that the social recognition deficit cannot be explained by a lack of sensitivity to social olfactory cues.

### 3.5. Effects of HFHS diet on novel object recognition

To confirm a role in cognition, we tested HFHS diet rats on their ability to explore novel compared to previously explored objects. Control rats showed intact object recognition memory by preferentially exploring the novel object; however, HFHS rats explored the familiar and novel objects equally, indicating impaired object recognition (object x diet group, *F*(1,14)=50.7, *P*<0.001; control *F*(1,14)=120.5, *P*<0.001, HFHS *F*<1, Fig 2H). Exploration ratios calculated from the test data [Mean (SEM): Control = 0.73 (0.01); HFHS = 0.52 (0.02)] differed significantly between groups (*F*(1,14)=60.8, *P*<0.001).

**Figure 2.**
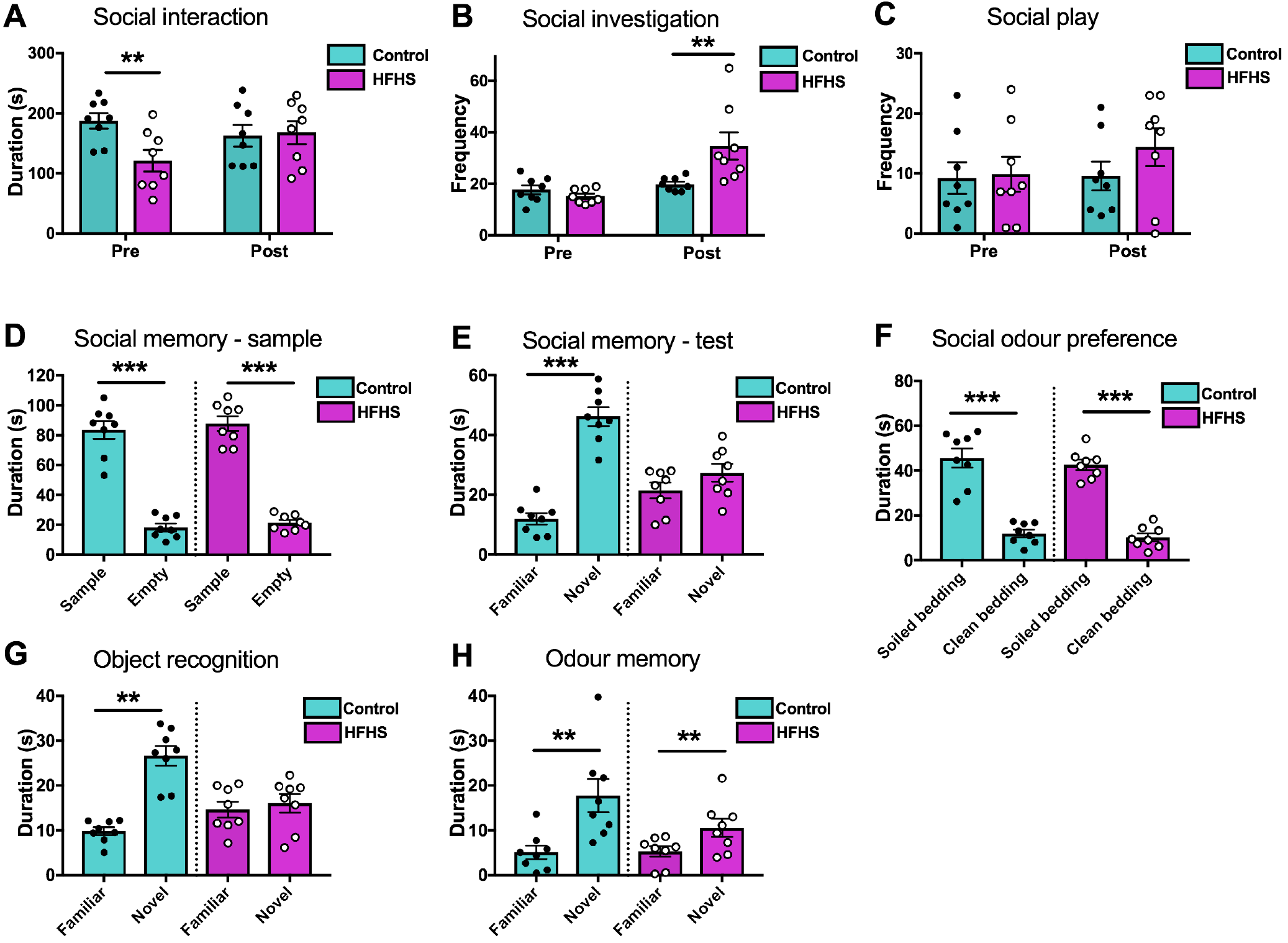
Social behaviours between control / HFHS diet exposed rats and a novel weight / age matched conspecific. HFHS diet rats were tested either 23h after HFHS pellet access “pre” or 1h after access to the HFHS pellets “post”, A) Total duration of social contact between rats, B) frequency of social interactions, and C) frequency of social play. Performance of HFHS diet and control rats in social recognition memory - D) exploration times of the chamber containing the sample rat “sample” and empty chamber “empty” during the sample phase of social memory testing, E) exploration times of the chamber containing the familiar sample rat and chamber containing a novel sample rat. F) Exploration time of chambers containing soiled bedding “social odour” or clean bedding. G) Novel odour recognition performance in control and HFHS diet rats during the test phase following a 5 min delay. H) Novel object recognition performance during the test phase following a 24h delay. Error bars represent +SEM. ** *P*<0.01. Error bars represent +SEM. * indicates *P*≤0.05, ** *P*<0.01, *** *P*<0.001 between groups comparisons.

### 3.6. Diet effects on PFC and hippocampal mRNA expression

To determine the whether short, intermittent periods of exposure to HFHS diet changed transcript expression within two brain regions associated with social behaviour, we quantified mRNA expression of genes related neuroplasticity, dopamine and monoamine signalling and inflammation (Table 1). We found the majority of transcript changes occurred in the prefrontal cortex. Consumption of the HFHS diet correlated with reduced expression of genes encoding enzymes involved in monoamine degradation, Comt and Maoa. The HFHS diet fed rats had reduced Maoa expression in the PFC (*F*(1,13)=8.50, *P*<0.05) and hippocampus (*F*(1,14)=6.89, *P*<0.05); Comt expression was reduced in the PFC (*F*(1,14)=19.0, *P*<0.001), but not the hippocampus. The neuroplasticity associated gene Bdnf was reduced in the PFC of HFHS consuming rats (*F*(1,13)=4.99, *P*<0.05). Group differences in PFC expression of the neuroinflammatory genes Nlrp3 (*F*(1,14)=4.41, *P*=0.056) and Il6 (*F*(1,14)=4.24, *P*=0.06) trended towards significance (Table 1).

**Table 1:** The effects of intermittent high fat and high sucrose (HFHS) diet exposure on prefrontal cortex and hippocampal gene expression. Table shows Mean (SEM), * = *P*<0.05, **=*P*<0.01, # = *P*<0.10

| Gene | Prefrontal cortex |  |  | Hippocampus |  |  |
| --- | --- | --- | --- | --- | --- | --- |
|  | Control | HFHS | *p*-value | Control | HFHS | *p*-value |
| **Neuroplasticity** |  |  |  |  |  |  |
| Gad1 | 0.84 (0.04) | 0.94 (0.07) | *n.s.* | 1.00 (0.08) | 0.94 (0.06) | *n.s.* |
| Bdnf | 1.00 (0.10) | 0.72 (0.03)* | 0.045 | 1.00 (0.08) | 0.96 (0.17) | *n.s.* |
| **Dopamine receptors** |  |  |  |  |  |  |
| Drd1a | 1.00 (0.23) | 0.65 (0.13) | *n.s.* | 1.00 (0.11) | 0.95 (0.09) | *n.s.* |
| Drd2 | 1.00 (0.32) | 0.56 (0.15) | *n.s.* | 0.93 (0.03) | 0.85 (0.07) | *n.s.* |
| **Monoamine synthesis** |  |  |  |  |  |  |
| Maoa | 1.00 (0.04) | 0.86 (0.02)* | 0.012 | 1.00 (0.04) | 0.83 (0.05)* | 0.02 |
| Comt | 1.00 (0.02) | 0.83 (0.03)** | 0.001 | 1.00 (0.07) | 1.08 (0.09) | *n.s.* |
| **Serotonin receptor** |  |  |  |  |  |  |
| Htr4 | 0.85 (0.07) | 0.79 (0.08) | *n.s.* | 1.00 (0.05) | 0.96 (0.04) | *n.s.* |
| **Inflammation** |  |  |  |  |  |  |
| Tnf-α | 1.00 (0.16) | 0.79 (0.11) | *n.s.* | 1.00 (0.21) | 0.65 (0.06) | *n.s.* |
| Il6 | 1.00 (0.15) | 0.62 (0.08)# | 0.060 | 0.81 (0.14) | 0.59 (0.11) | *n.s.* |
| Nlrp3 | 1.00 (0.06) | 0.85 (0.03)# | 0.056 | 1.00 (0.18) | 1.03 (0.14) | *n.s.* |
| Itgam | 1.00 (0.10) | 1.09 (0.09) | *n.s.* | 1.00 (0.11) | 1.23 (0.19) | *n.s.* |

### 3.7. Microbiota composition and analysis

The relative abundance of a number of specific taxa differed significantly between the two diet groups as shown by DESEq2 analysis (Fig 3A, Supplementary Table 3). HFHS diet increased bacteria from *Firmicutes* phylum *Clostridales* family, including *Lachnospiraceae* (Genus *Blauta*, q<0.04; Genus Unspecified q<0.03), *Ruminoccoceae* (Genus Unspecified q<0.01) and *Veillonellaceae* (Genus *Phascolarctobacterium* q<0.02). HFHS diet increased bacteria from *Actinobacteria* phylum, family *Bifidobacteriaceae* (Genus *Bifidobacterium* q<0.04), *Bacteroidetes* phylum, order *Bacteroidales* (Genus Unspecified q<0.05) and *Tenericutes* phylum, order *Erysipelotrichaceae* (Genus *Allobaculum* q<0.05).

Unsupervised principle component analysis (PCA) revealed overlap between groups, the first component explained 22% of variance; the second component 11% (Fig 3B). Partial least squares discriminant analysis (PLS-DA) identified the two components that discriminate maximally between the HFHS and control diet groups, showing a large proportion of variance accounted for by the first component (21%) and a lesser degree by the second (8%), this demonstrated significant separation of the microbiota community structure between the groups (Fig 3C). Alpha diversity did not differ between the HFHS and control groups measured by observed species, Chao 1, Shannon or Simpson indices (see Fig 3D; *Fs*<1).

**Figure 3.**
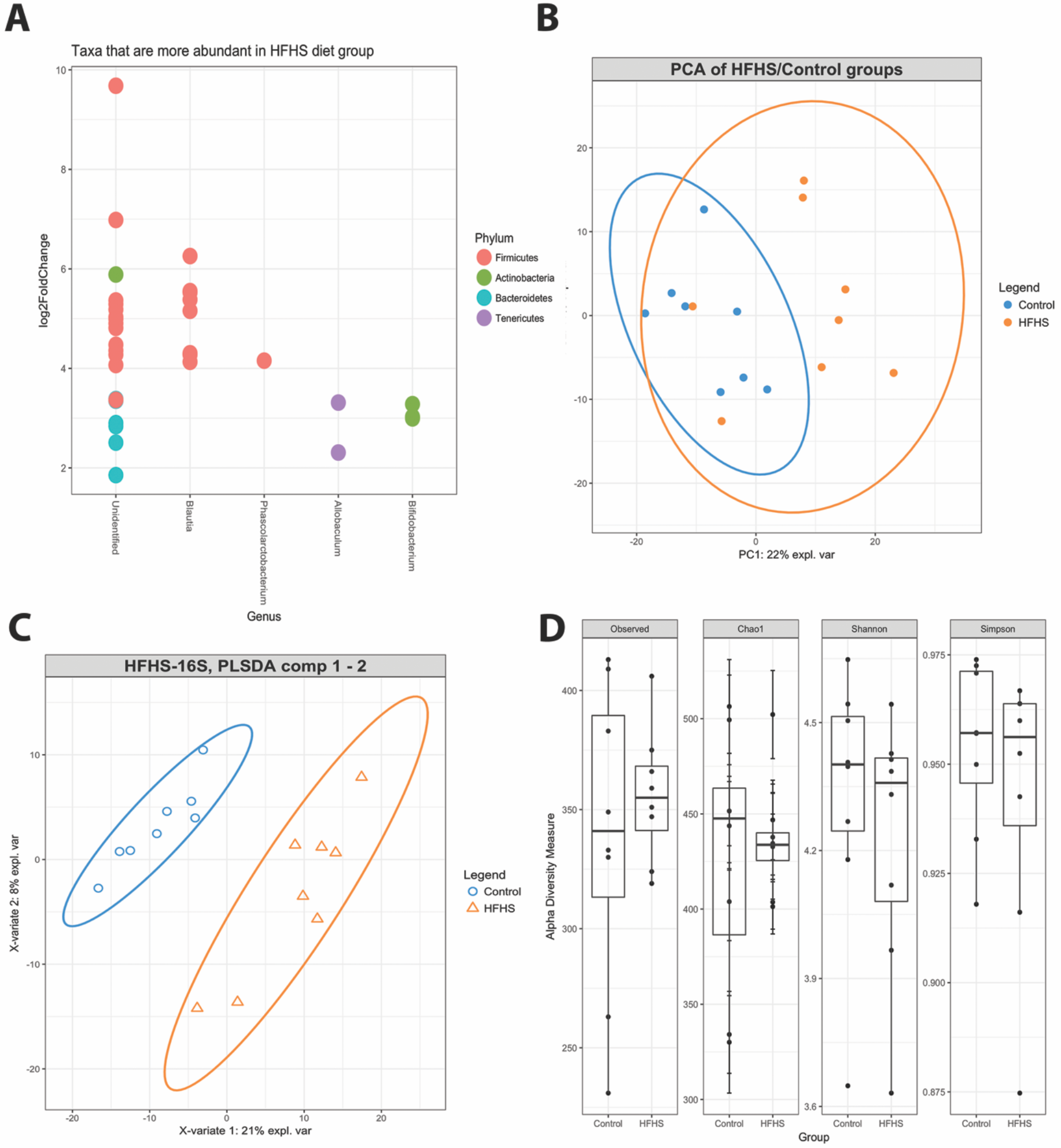
A) Graphical depiction of DESEq2 analysis. Each coloured circle represents one bacterial genus that was more abundant in the HFHS than control group (q <0.05). Log2 fold change refers to the difference abundance of the log2 values between diet groups for each bacterial genus. B) Unsupervised Principle Component Analysis (PCA) plot of microbiome samples from the HFHS diet (orange dots) and control diet (blue dots). The first component explained 22% of variance; the second component 11%. C) Partial least-squares discriminant analysis (PLS-DA) figure showing a large proportion of variance accounted for by the first component (21%) and a lesser degree by the second (8%). Each point represents a sample. D) No significant differences in alpha diversity of faecal microbiota between HFHS and control diet groups. Each panel represents one alpha diversity measure as follows: Observed = total number of OTU’s observed; Chao1 = richness estimator (estimate of the total number of OTU’s present in a community); Shannon and Simpson = microbial indexes of diversity. Boxes span the first to third quartiles; the horizontal line inside the boxes represents the median. Whiskers extending vertically from the boxes indicate variability outside the upper and lower quartiles, and the single black circles indicate outliers (all *P*>0.05).

### 3.8. Associations between diet effects, behavioural performance and gene expression

Correlations were performed between individual representative values of specific behaviours (social interaction pre consumption of diet, social recognition and novel object recognition) that differed between diet group and biological measurements (adiposity and cortical gene expression).

A number of associations were observed, in particular positive correlations between PFC expression of Maoa and social interaction pre-HFHS diet and object memory. Full bivariate correlation analysis presented in Figure 4.

**Figure 4.**
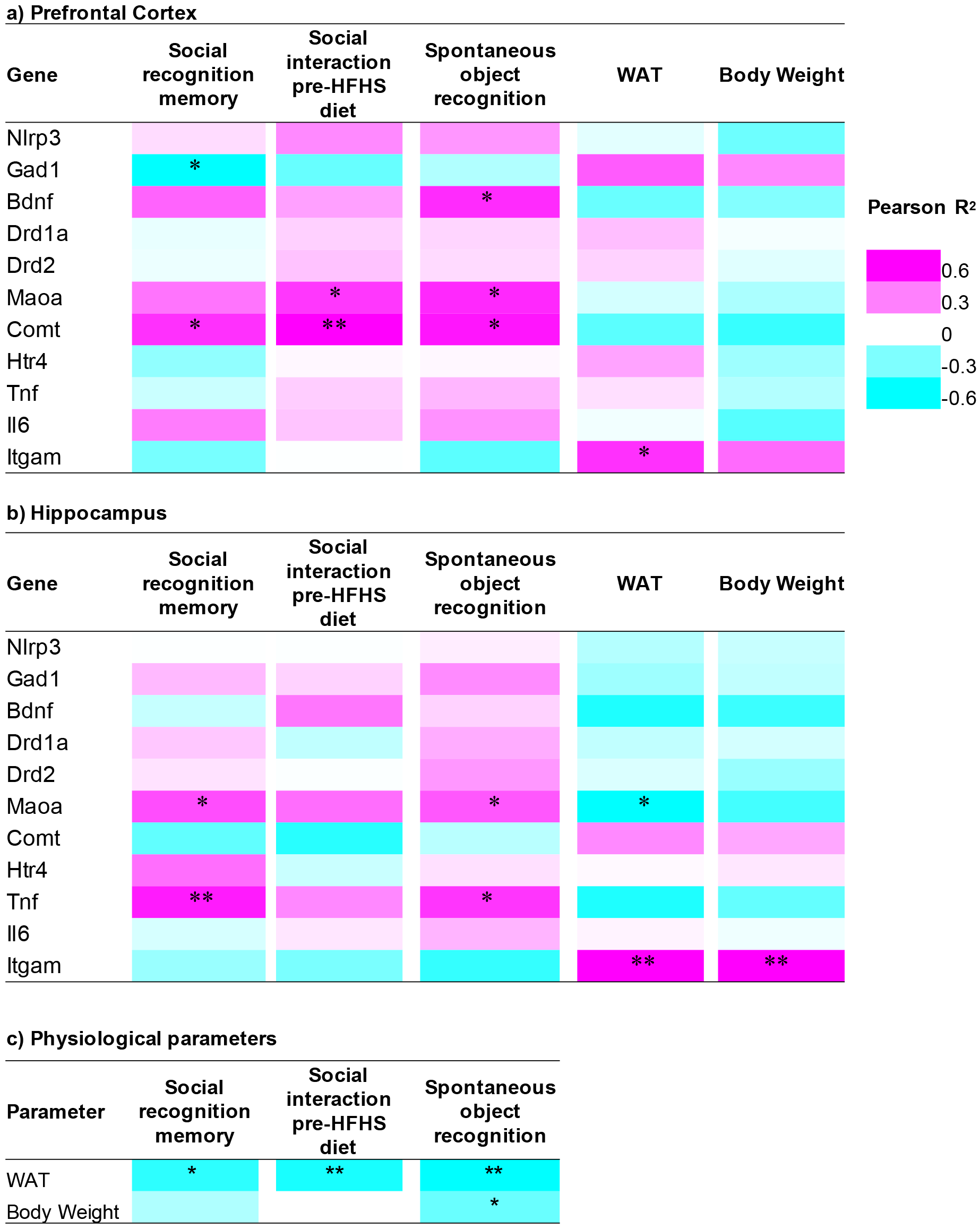
Heatmap of bivariate correlations (Pearson’s R^2^) between behavioural assays – social recognition memory, social interaction and novel object recognition performance, and a) prefrontal cortex gene expression, b) hippocampal gene expression and c) physiological parameters. *=*P*<0.05, **=*P*<0.01

A number of bivariate correlations between physiological parameters (WAT and bodyweight) and gene expression were significant (Figure 4a and b). In particular, PFC and hippocampal Itgam expression was positively correlated with WAT (PFC: r^2^=0.52, *P*<0.05, HPC: r^2^=0.66, *P*<0.01) and bodyweight (HPC: r^2^=0.67, *P*<0.01), and hippocampal Maoa expression was negatively correlated with WAT (r^2^=−0.45, *P*<0.05). Correlations between physiological parameters (WAT and bodyweight) and behavioural performance were observed (Figure 4c), in particular significant negative correlations between WAT and social recognition memory (r^2^=<0.56, *P*<0.05), social interaction pre-HFHS diet (r^2^=−0.58, *P*<0.01) and novel object recognition performance (r^2^=−0.65, *P*<0.01).

Total WAT was significantly associated with PFC gene expression (*F*(1,12)=5.4, *P*<0.05); specifically Tnf-α (adjusted r^2^=0.41, *P*<0.01), Comt (adjusted r^2^=0.23, *P*<0.05), Maoa (adjusted r^2^=0.29, *P*<0.05), and Bdnf (adjusted r^2^=0.74, *P*<0.001). A number of bivariate correlations between bodyweight and gene expression were significant (Figure 4a and b) however these associations did not persist in multivariate linear modelling (overall model *F*(1,12)=2.1, *P*=0.17). There were no significant associations between hippocampal gene expression and body weight (*F*(1,13)<1). WAT weight predicted Il6 expression in the hippocampus (*F*(1,13)=4.86, *P*=0.05).

Associations between hippocampal and PFC genes differentially expressed in control and HFHS groups (see Table 1, Figure 4) and behavioural performance were examined. No predictive relationships were observed between PFC Bdnf, Comt or Maoa expression and social interaction pre diet consumption, social memory or novel object recognition (*P*=0.17; *P*=0.092; *P*=0.16 for overall model of each gene respectively). There was no evidence for a predictive relationship between hippocampal Maoa expression and behaviours (*P*=0.35).

### 3.9. Associations between gut microbiota composition and social behaviour

Scores on pre-diet social behaviour, social recognition memory and novel object recognition tasks respectively were all significantly associated with the relative abundance of a number of bacterial taxa (all associations where q<0.05 presented in Table 2).

Social memory performance was associated with a large number of taxa. Higher social memory scores were associated with a greater abundance of bacteria from the *Bifidobacteriales* and *Bacteroidales* order, *Lachnospiraceae* family (*Blautia* and multiple unspecified genera), *Ruminococcaceae* family and genus *Allobaculum*. Novel object recognition was negatively associated with abundance of *Bacteroidales* and a number of taxa from the *Lachnospiraceae* family. Only three taxa were significantly associated with social behaviour pre HFHS diet: a relative reduction of *Bifidobacteriales* order and two unspecified genera from the *Lachnospiraceae* family.

**Table 2.**
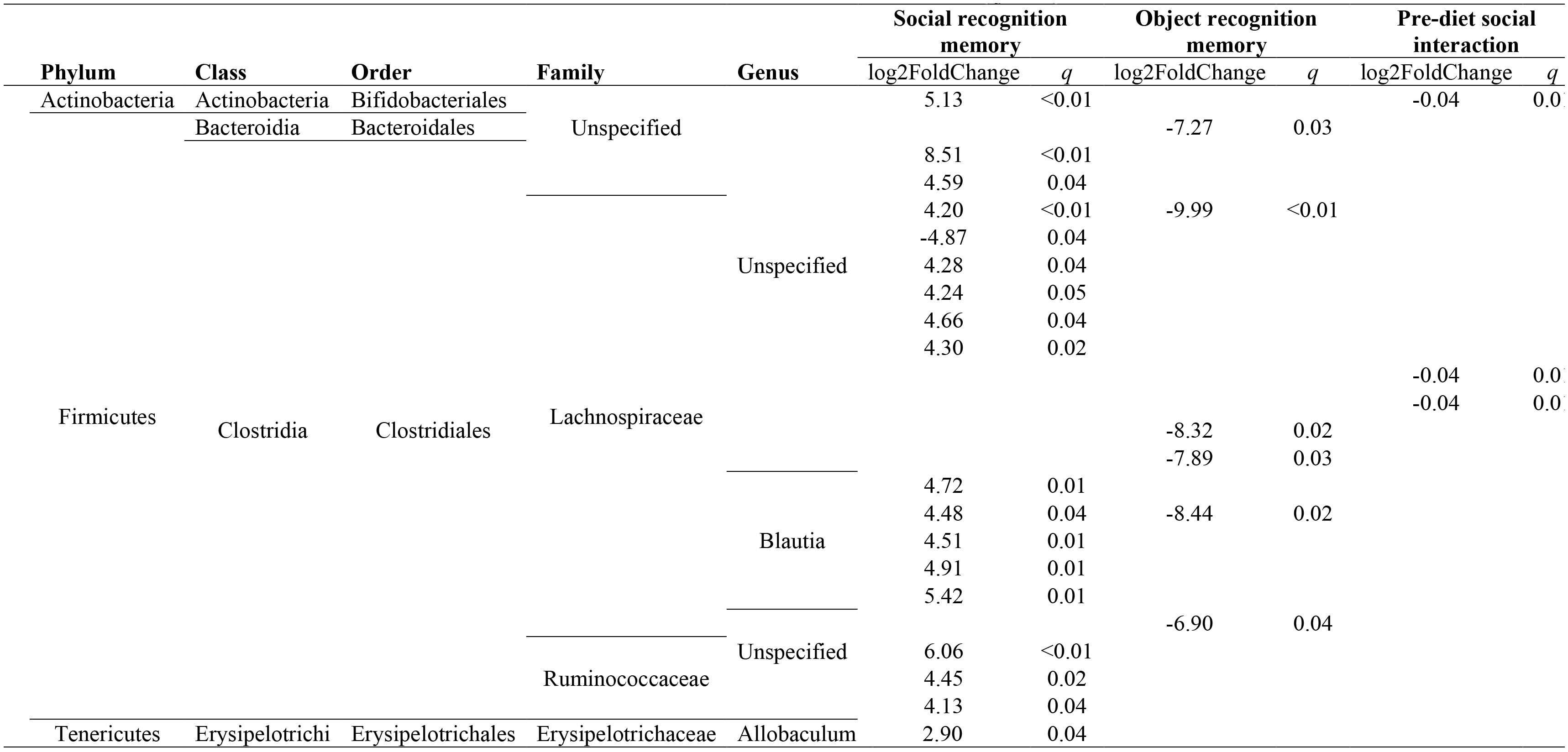
Associations between relative abundance of taxa in faecal microbiota and behavioural outcomes (q<0.05)

### 3.10. Firmicutes to Bacteroidetes ratio

There were no significant differences between the diet groups on *Firmicutes* to *Bacteroidetes* ratio (FB ratio; t(9.63)=−1.03, *P*=0.33). Samples were pooled for subsequent FB ratio analyses, with diet group included to control for potential interaction effects. Multivariate linear modelling demonstrated a significant relationship between FB ratio and the three behavioural dependent variables: social memory, novel object recognition and pre-diet social interaction (*F*(3,11)=5.26, *P*=0.02). Post-hoc tests demonstrated strong evidence that FB ratio negatively predicted “pre” diet social behaviour (F(2,13)=11.46, *P*=0.001), but not object or social recognition memory.

### 3.11. Associations between gut microbiota and hippocampal /PFC gene expression

The hippocampal and PFC genes found to differ in expression between the control and HFHS diet groups (PFC: Bdnf, Maoa, Comt; hippocampus: Maoa, *P*s<0.05) were tested for their associations with differential abundance of bacterial taxa. Of these, significantly differentially abundant taxa (q<0.05) were apparent only for Maoa (Table 3). PFC Maoa expression was positively associated with one genera of the *Lachnospiraceae* family, whilst a number of bacteria across the four primary phyla were differentially abundant on the basis of hippocampal Maoa expression in both positive and negative directions.

**Table 3.**
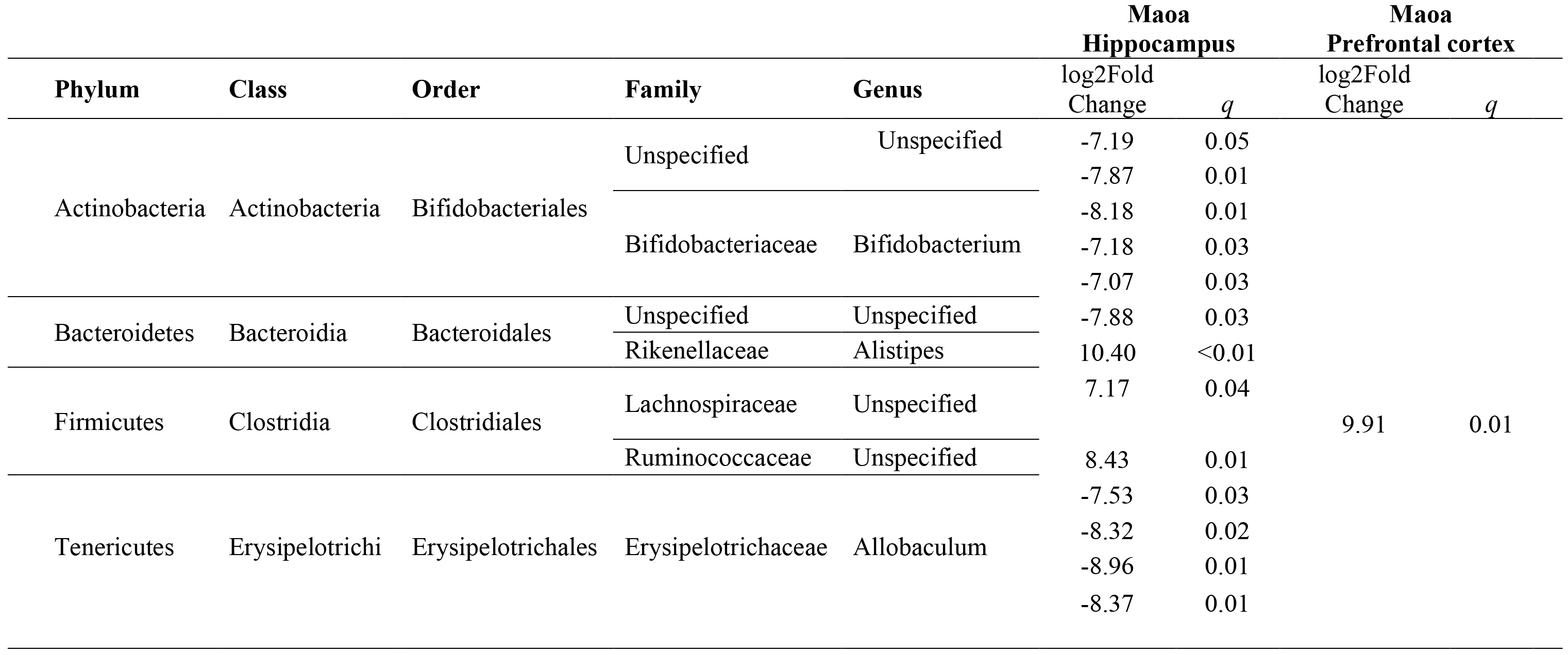
Associations between relative abundance of taxa in faecal microbiota and gene expression (q<0.05).

## 4. Discussion

Here we show that daily limited consumption of a HFHS diet leads to alterations in social interaction and social memory in young rats, and impaired novel object recognition memory. Previous studies have shown that high-energy diets rapidly cause hippocampal-dependent memory deficits (Kanoski et al., 2007), but none have examined the impact of intermittent HFHS diets on social behaviours in rats. This is the first demonstration to associate diet-induced alterations to social behaviour with microbiota and brain changes in reward neurotransmission and neuroplasticity.

Intermittent access to a HFHS diet influenced the presentation of normal social behaviours, including social interaction and preference for social novelty. Consumption of a HFHS diet for 2h/day reduced duration engaged in social interaction prior to diet availability, but not following diet consumption. This indicates that when rats expected to receive the HFHS diet they were less willing to engage in social interaction, potentially underpinned by diet evoked alterations in Maoa and Comt expression, leading to changes in monoamine neurotransmission, or increased anxiety. Thus, limited access to a HFHS diet may influence social interaction, as comparable interaction durations were observed following access to the HFHS food. Moreover, social interaction frequency was increased after rats had access to the HFHS food, suggesting that following consumption of HFHS diet these rats may find interaction more rewarding, or may have reduced anxiety. Social play is important for neurobehavioural development; however, we did not observe differences in frequencies between diet groups. This may be due to the group housing conditions and short period of isolation used prior to behavioural testing; as a recent study reported that isolation amplified social play behaviour (Carvalho et al., 2016). Another possible explanation is that social play declines across adolescence, and that the apparent lack of social play differences was due to the rats age at testing (mid-late adolescence) (Trezza et al., 2010). Further studies should be conducted to examine whether diet manipulations specifically during adolescence endure into adulthood to identify whether adolescence poses as a critical window of vulnerability to social behavioural changes.

Social recognition memory differed between control and HFHS rats, with HFHS rats showing no observed preference for the chamber containing the novel rat during the test phase, indicative of impaired social memory. This complements a recent study showing that an acute exposure to high fat diet in juvenile rats impaired social memory (Yaseen et al., 2018). As rats showed differences in their duration of time engaged in social interaction prior to consuming the HFHS food, the social memory testing was conducted following HFHS access, to ensure that any memory deficits observed were not due to reduced social contact in the HFHS diet rats. Initial sociability during the sample phase did not differ between HFHS and control diet rats, thus it appears that social memory was impacted specifically by diet. Social memory has been shown to depend upon both PFC and hippocampal function (Kogan et al., 2000; Tanimizu et al., 2017), and our observed alterations to markers of monoamine neurotransmission and neuroplasticity may underpin social changes. This is also complemented by impaired long term novel object recognition, again indicative of hippocampal dysfunction (Warburton and Brown, 2010). Moreover, both HFHS and control diet rats showed preference for a social odour and showed intact odour recognition memory. Thus, intermittent HFHS diet did not impact olfactory discrimination, and highlights that social memory deficits are unlikely to be underpinned by impaired odour discrimination.

Diet-correlated alterations to mRNA expression of enzymes Maoa and Comt, were observed in the PFC, indicating that HFHS diet consumption impacts on monoamine neurotransmission integral for social behaviour and cognition. Dopamine is a critical neurotransmitter in the regulation of food intake; in particular dopamine activity in the mesocorticolimbic dopamine circuitry is associated with food reward (Volkow et al., 2011). Maoa, the gene for monoamine oxidase, deaminates dopamine and has a key role in controlling the availability of cortical dopamine. Similarly, Comt is involved in the degradation of dopamine. Changes to monoamine signalling may therefore underpin the altered social behaviour and social memory observed in HFHS diet rats, supported by reports of diet-induced alterations to dopamine receptor expression in the striatum (Johnson and Kenny, 2010). However, we observed no dopamine receptor (Drd1a/Drd2) expression changes in the hippocampus or PFC, suggesting that these receptor mRNA changes may be specific to striatal regions following palatable high sugar diets (Naneix et al., 2018; Naneix et al., 2017). Further studies should examine whether other reward associated genes, such as serotonin and mu-opioid receptors are altered by this diet protocol, and also the involvement of oxytocin signalling mechanisms (Yaseen et al., 2018).

Reduced PFC Bdnf expression was observed in HFHS consuming rats, which also correlated positively with novel object recognition performance. This diet-induced change may underpin the changes to social behaviours and cognition as BDNF signalling has a critical role in memory encoding (Choi et al., 2010). Decreased levels of BDNF in the hypothalamus, PFC or serum have been shown to correlate with depression-like behaviours in animals and humans (Bocchio-Chiavetto et al., 2010) and high fat diet consumption reduces hippocampal BDNF levels (Molteni et al., 2004; Pistell et al., 2010) linking BDNF to emotional processes. Gut microbiota composition is also linked to alterations in BDNF within regions essential for learning and emotional behaviours, as demonstrated by previous studies indicate reduced cortical and hippocampal Bdnf gene expression in GF mice (Sudo et al., 2004), and antibiotic-induced microbiota dysbiosis altered protein levels of BDNF in the amygdala and hippocampus as well as reduced anxiety-like behaviours in the light-dark box (Bercik et al., 2011). This provides a mechanistic insight into the influence of the microbiome in cognition and emotional regulation via BDNF expression.

Excessive consumption of saturated fats is shown to induce secretion of pro-inflammatory cytokines by adipocytes and macrophages, and affect the integrity of the blood-brain barrier (BBB) (Kanoski et al., 2010), allowing pro-inflammatory cytokines and immune cells to reach the brain (Thaler et al., 2012). Interestingly, no significant changes between groups were observed to inflammatory marker mRNA expression (Il6, TNF-α, Nlrp3, Itgam) were observed in this study, and trends indicated that PFC expression of Il6 and Nlrp3 was lower in HFHS diet rats compared to controls. This may be due to the age of the rats, as emergent evidence suggests that the modulatory effects of obesogenic diets on inflammatory markers occur in an age-dependent manner, with younger rats showing resistance to neuroinflammation (Teixeira et al., 2017). However, Itgam (cluster of differentiation molecule 11b, CD11b) expression in the PFC and hippocampus positively correlated with WAT, indicative that increased adiposity was associated with aspects of neuroinflammation. Itgam is expressed by microglia, and also neutrophils and monocytes in the injured brain (Jeong et al., 2013). More so, evidence indicates that obesity-induced neuroinflammation is dependent on the type of diet in terms of fat and sugar content, the duration of the diet and regional differences in brain structures (Guillemot-Legris and Muccioli, 2017). Future studies utilising immunohistochemistry to examine microglia morphology and astrogliosis and protein markers are needed to validate the region specific impact of obesogenic diets on mPFC and hippocampal neuroinflammatory markers.

HFHS diet consumption resulted in significantly increased WAT deposits characteristic of diet-induced obesity, and has been previously associated with decreased abundance of *Bacteroidetes* and increases in *Firmicutes* bacteria (Ley et al., 2005); however, we did not observe overall alterations to the *Firmicutes* to *Bacteriodetes* ratio with this intermittent feeding schedule. Our study suggests that members of *Bacteroites* (order *Bacteroidales)* were significantly increased in rats that consumed HFHS diets. Therefore, not all the members of the *Bacteriodetes* family are decreased with adiposity. However, it is possible that *Firmicutes* to *Bacteriodetes* ratio changes become more prominent with the ongoing development of obesity and chronic consumption of HFHS diets, rather than short-term intake of such foods. Differential abundance analyses demonstrated that taxa from *Lachnospiraceae* and *Ruminoccoceae* families of the *Clostridiales* order were the most common bacterial predictors of social behaviour and recognition memory, converging with clinical studies that show similar changes to microbiome populations in neuropsychiatric disorders including major depressive disorder and autism (De Angelis et al., 2013; Naseribafrouei et al., 2014). Moreover, social avoidance behaviour in adult mice is associated with increased abundance of *Lachnospiraceae*, *Ruminococcaceae* and *Clostridiales* in non-obese diabetic mice, and the transfer of intestinal microbiota from these mice to microbiota-depleted recipients evoked similar behavioural phenotypes (Gacias et al., 2016), suggesting a key role of diet and metabolism in microbiota signatures. As such, converging evidence indicates the potential influence of diet on social development and social behaviours via the gut-brain-microbiota axis (Christian et al., 2015; Parashar and Udayabanu, 2016).

### Future studies

Future studies should examine whether faecal transplants from HFHS diet animals evokes similar behavioural and cortical gene expression changes in Maoa, Comt and Bdnf allowing for increased mechanistic insights into the effects of the microbiome on the brain. Furthermore, predictions of the metagenome functional content from the bacterial communities would further provide insight into the metabolic pathways affected by intermittent HFHS diet consumption. Modulation of the gut-brain axis dynamics has clinical implications for mental health conditions, and as such the use of “psychobiotics” is posited as a novel therapeutic avenue for psychiatric disorders and diet-induced cognitive changes. Treatment strategies that target the gut microbiome should be explored, such as commensal bacteria, which have been shown to ameliorate depressive (Bravo et al., 2011) and anxiety like behaviour (Bercik et al., 2011), and prebiotics which increase Bdnf mRNA expression in the hippocampus (Burokas et al., 2017). These may provide a route for the attenuation of diet and obesity evoked cognitive and emotional alterations.

### Conclusions and implications

In conclusion, these results demonstrate that intermittent access to a HFHS diet can rapidly impact social behaviour and cognition, evoke alterations in cortical gene expression, and alter microbiota composition. Modulation of the microbiota may lead to the emergence of novel therapies to combat social, emotional and cognitive deficits, which have been linked with metabolic disorders, and for the treatment of neuropsychiatric disorders.

## List of abbreviations

ACTB: Actin, beta
ANOVA: Analysis of variance
BDNF: Brain derived neurotrophic factor
CD11b: cluster of differentiation molecule 11b
Comt: catechol-O-methyltransferase
DRD1: Dopamine receptor D1
DRD2: Dopamine receptor D2
FB ratio: *Firmicutes* to *Bacteroidetes* ratio
GAD1: Glutamate decarboxylase 1
GF: Germ free
gnWAT: Gonadal white adipose tissues HFHS - High fat and high sugar
HTR4: 5-hydroxytryptamine (serotonin) receptor 4, G-coupled
IL6: Interleukin 6
ITGAM: Integrin, alpha M
PFC: Prefrontal cortex
MANOVA: Multivariate analysis of variance
Maoa: Monoamine Oxidase
mRNA: messenger ribonucleic acid
Nlrp3: NLR family, pyrin domain containing 3
OTU: Operational taxonomic units
P: Postnatal day
PCA: Principle Component Analysis
PLS-DA: Partial least-squares discriminant analysis
QIIME: Quantitative Insights Into Microbial Ecology
rpWAT: Retroperitoneal white adipose tissues
Tnf-α: Tumour necrosis factor alpha

## 5. Acknowledgments

This work was supported by an Australian Research Council Discovery Early Career Research Award (DE140101071) to ACR. JD receives funding from the Royal Society (UK) grant (RG130316) and an Alzheimer’s Society Fellowship (AS-JF-15-008). The authors wish to thank Professor Anthony Hannan for his helpful feedback on the manuscript. All authors report no biomedical financial interests or potential conflicts of interest.

## References

Albenberg, L. G., Wu, G. D., 2014. Diet and the intestinal microbiome: associations, functions, and implications for health and disease. Gastroenterology 146, 1564–1572.

Ashelford, K. E., Chuzhanova, N. A., Fry, J. C., Jones, A. J., Weightman, A. J., 2005. At least 1 in 20 16S rRNA sequence records currently held in public repositories is estimated to contain substantial anomalies. Appl Environ Microbiol 71, 7724–7736.

Baker, K. D., Loughman, A., Spencer, S. J., Reichelt, A. C., 2017. The impact of obesity and hypercaloric diet consumption on anxiety and emotional behavior across the lifespan. Neurosci Biobehav Rev 83, 173–182.

Baker, K. D., Reichelt, A. C., 2016. Impaired fear extinction retention and increased anxiety like behaviours induced by limited daily access to a high-fat/high-sugar diet in male rats: Implications for diet-induced prefrontal cortex dysregulation. Neurobiol Learn Mem 136, 127–138.

Benjamini, Y., Drai, D., Elmer, G., Kafkafi, N., Golani, I., 2001. Controlling the false discovery rate in behavior genetics research. Behav Brain Res 125, 279–284.

Bercik, P., Denou, E., Collins, J., Jackson, W., Lu, J., Jury, J., Deng, Y., Blennerhassett, P., Macri, J., McCoy, K. D., Verdu, E. F., Collins, S. M., 2011. The intestinal microbiota affect central levels of brain-derived neurotropic factor and behavior in mice. Gastroenterology 141, 599-609, 609 e591–593.

Bicks, L. K., Koike, H., Akbarian, S., Morishita, H., 2015. Prefrontal Cortex and Social Cognition in Mouse and Man. Front Psychol 6, 1805.

Bocarsly, M. E., Hoebel, B. G., Paredes, D., von Loga, I., Murray, S. M., Wang, M., Arolfo, M. P., Yao, L., Diamond, I., Avena, N. M., 2014. GS 455534 selectively suppresses binge eating of palatable food and attenuates dopamine release in the accumbens of sugar-bingeing rats. Behav Pharmacol 25, 147–157.

Bocchio-Chiavetto, L., Bagnardi, V., Zanardini, R., Molteni, R., Nielsen, M. G., Placentino, A., Giovannini, C., Rillosi, L., Ventriglia, M., Riva, M. A., Gennarelli, M., 2010. Serum and plasma BDNF levels in major depression: a replication study and meta-analyses. World J Biol Psychiatry 11, 763–773.

Braithwaite, I., Stewart, A. W., Hancox, R. J., Beasley, R., Murphy, R., Mitchell, E. A., Group, I. P. T. S., Group, I. P. T. S., 2014. Fast-food consumption and body mass index in children and adolescents: an international cross-sectional study. BMJ Open 4, e005813.

Bravo, J. A., Dinan, T. G., Cryan, J. F., 2011. Alterations in the central CRF system of two different rat models of comorbid depression and functional gastrointestinal disorders. Int J Neuropsychopharmacol 14, 666–683.

Burokas, A., Arboleya, S., Moloney, R. D., Peterson, V. L., Murphy, K., Clarke, G., Stanton, C., Dinan, T. G., Cryan, J. F., 2017. Targeting the Microbiota-Gut-Brain Axis: Prebiotics Have Anxiolytic and Antidepressant-like Effects and Reverse the Impact of Chronic Stress in Mice. Biol Psychiatry 82, 472–487.

Carvalho, A. L. O., Ferri, B. G., de Sousa, F. A. L., Vilela, F. C., Giusti-Paiva, A., 2016. Early life overnutrition induced by litter size manipulation decreases social play behavior in adolescent male rats. Int J Dev Neurosci 53, 75–82.

Choi, D. C., Maguschak, K. A., Ye, K., Jang, S. W., Myers, K. M., Ressler, K. J., 2010. Prelimbic cortical BDNF is required for memory of learned fear but not extinction or innate fear. Proc Natl Acad Sci U S A 107, 2675–2680.

Christian, L. M., Galley, J. D., Hade, E. M., Schoppe-Sullivan, S., Kamp Dush, C., Bailey, M. T., 2015. Gut microbiome composition is associated with temperament during early childhood. Brain Behav Immun 45, 118–127.

Crawley, J. N., Chen, T., Puri, A., Washburn, R., Sullivan, T. L., Hill, J. M., Young, N. B., Nadler, J. J., Moy, S. S., Young, L. J., Caldwell, H. K., Young, W. S., 2007. Social approach behaviors in oxytocin knockout mice: comparison of two independent lines tested in different laboratory environments. Neuropeptides 41, 145–163.

Davis, J. F., Tracy, A. L., Schurdak, J. D., Tschop, M. H., Lipton, J. W., Clegg, D. J., Benoit, S. C., 2008. Exposure to elevated levels of dietary fat attenuates psychostimulant reward and mesolimbic dopamine turnover in the rat. Behav Neurosci 122, 1257–1263.

De Angelis, M., Piccolo, M., Vannini, L., Siragusa, S., De Giacomo, A., Serrazzanetti, D. I., Cristofori, F., Guerzoni, M. E., Gobbetti, M., Francavilla, R., 2013. Fecal microbiota and metabolome of children with autism and pervasive developmental disorder not otherwise specified. PLoS One 8, e76993.

Del Rio, D., Morales, L., Ruiz-Gayo, M., Del Olmo, N., 2016. Effect of high-fat diets on mood and learning performance in adolescent mice. Behav Brain Res 311, 167–172.

DeSantis, T. Z., Hugenholtz, P., Larsen, N., Rojas, M., Brodie, E. L., Keller, K., Huber, T., Dalevi, D., Hu, P., Andersen, G. L., 2006. Greengenes, a chimera-checked 16S rRNA gene database and workbench compatible with ARB. Appl Environ Microbiol 72,5069–5072.

Desbonnet, L., Clarke, G., Traplin, A., O’Sullivan, O., Crispie, F., Moloney, R. D., Cotter, P. D., Dinan, T. G., Cryan, J. F., 2015. Gut microbiota depletion from early adolescence in mice: Implications for brain and behaviour. Brain Behav Immun 48, 165–173.

Edgar, R. C., 2010. Search and clustering orders of magnitude faster than BLAST. Bioinformatics 26, 2460–2461.

Fadrosh, D. W., Ma, B., Gajer, P., Sengamalay, N., Ott, S., Brotman, R. M., Ravel, J., 2014. An improved dual-indexing approach for multiplexed 16S rRNA gene sequencing on the Illumina MiSeq platform. Microbiome 2, 6.

Frohlich, E. E., Farzi, A., Mayerhofer, R., Reichmann, F., Jacan, A., Wagner, B., Zinser, E., Bordag, N., Magnes, C., Frohlich, E., Kashofer, K., Gorkiewicz, G., Holzer, P., 2016. Cognitive impairment by antibiotic-induced gut dysbiosis: Analysis of gut microbiota-brain communication. Brain Behav Immun 56, 140–155.

Furlong, T. M., Jayaweera, H. K., Balleine, B. W., Corbit, L. H., 2014. Binge-like consumption of a palatable food accelerates habitual control of behavior and is dependent on activation of the dorsolateral striatum. J Neurosci 34, 5012–5022.

Gacias, M., Gaspari, S., Santos, P. M., Tamburini, S., Andrade, M., Zhang, F., Shen, N., Tolstikov, V., Kiebish, M. A., Dupree, J. L., Zachariou, V., Clemente, J. C., Casaccia, P., 2016. Microbiota-driven transcriptional changes in prefrontal cortex override genetic differences in social behavior. Elife 5.

Guillemot-Legris, O., Muccioli, G. G., 2017. Obesity-Induced Neuroinflammation: Beyond the Hypothalamus. Trends Neurosci 40, 237–253.

Jeong, H. K., Ji, K., Min, K., Joe, E. H., 2013. Brain inflammation and microglia: facts and misconceptions. Exp Neurobiol 22, 59–67.

Johnson, P. M., Kenny, P. J., 2010. Dopamine D2 receptors in addiction-like reward dysfunction and compulsive eating in obese rats. Nat Neurosci 13, 635–641.

Kanoski, S. E., Meisel, R. L., Mullins, A. J., Davidson, T. L., 2007. The effects of energy-rich diets on discrimination reversal learning and on BDNF in the hippocampus and prefrontal cortex of the rat. Behav Brain Res 182, 57–66.

Kanoski, S. E., Zhang, Y., Zheng, W., Davidson, T. L., 2010. The effects of a high-energy diet on hippocampal function and blood-brain barrier integrity in the rat. J Alzheimers Dis 21, 207–219.

Kim, Y., Venkataraju, K. U., Pradhan, K., Mende, C., Taranda, J., Turaga, S. C., Arganda-Carreras, I., Ng, L., Hawrylycz, M. J., Rockland, K. S., Seung, H. S., Osten, P., 2015. Mapping social behavior-induced brain activation at cellular resolution in the mouse. Cell Rep 10, 292–305.

Kogan, J. H., Frankland, P. W., Silva, A. J., 2000. Long-term memory underlying hippocampus-dependent social recognition in mice. Hippocampus 10,47–56.

Kolb, B., 1974. Social behavior of rats with chronic prefrontal lesions. J Comp Physiol Psychol 87, 466–474.

Labouesse, M. A., Lassalle, O., Richetto, J., Iafrati, J., Weber-Stadlbauer, U., Notter, T., Gschwind, T., Pujadas, L., Soriano, E., Reichelt, A. C., Labouesse, C., Langhans, W., Chavis, P., Meyer, U., 2017. Hypervulnerability of the adolescent prefrontal cortex to nutritional stress via reelin deficiency. Mol Psychiatry 22, 961–971.

Ley, R. E., Backhed, F., Turnbaugh, P., Lozupone, C. A., Knight, R. D., Gordon, J. I., 2005. Obesity alters gut microbial ecology. Proc Natl Acad Sci U S A 102, 11070–11075.

Livak, K. J., Schmittgen, T. D., 2001. Analysis of relative gene expression data using realtime quantitative PCR and the 2(-Delta Delta C(T)) Method. Methods 25, 402–408.

Love, M. I., Huber, W., Anders, S., 2014. Moderated estimation of fold change and dispersion for RNA-seq data with DESeq2. Genome Biol 15, 550.

Molteni, R., Wu, A., Vaynman, S., Ying, Z., Barnard, R. J., Gomez-Pinilla, F., 2004.Exercise reverses the harmful effects of consumption of a high-fat diet on synaptic and behavioral plasticity associated to the action of brain-derived neurotrophic factor. Neuroscience 123, 429–440.

Naneix, F., Darlot, F., De Smedt-Peyrusse, V., Pape, J. R., Coutureau, E., Cador, M., 2018. Protracted motivational dopamine-related deficits following adolescence sugar overconsumption. Neuropharmacology 129, 16–25.

Naneix, F., Tantot, F., Glangetas, C., Kaufling, J., Janthakhin, Y., Boitard, C., De Smedt-Peyrusse, V., Pape, J. R., Vancassel, S., Trifilieff, P., Georges, F., Coutureau, E., Ferreira, G., 2017. Impact of Early Consumption of High-Fat Diet on the Mesolimbic Dopaminergic System. eNeuro 4.

Naseribafrouei, A., Hestad, K., Avershina, E., Sekelja, M., Linlokken, A., Wilson, R., Rudi, K., 2014. Correlation between the human fecal microbiota and depression. Neurogastroenterol Motil 26, 1155–1162.

Ogden, C. L., Carroll, M. D., Kit, B. K., Flegal, K. M., 2014. Prevalence of childhood and adult obesity in the United States, 2011-2012. JAMA 311, 806–814.

Okuyama, T., Kitamura, T., Roy, D. S., Itohara, S., Tonegawa, S., 2016. Ventral CA1 neurons store social memory. Science 353, 1536–1541.

Parashar, A., Udayabanu, M., 2016. Gut microbiota regulates key modulators of social behavior. Eur Neuropsychopharmacol 26, 78–91.

Pistell, P. J., Morrison, C. D., Gupta, S., Knight, A. G., Keller, J. N., Ingram, D. K., Bruce-Keller, A. J., 2010. Cognitive impairment following high fat diet consumption is associated with brain inflammation. J Neuroimmunol 219, 25–32.

Reichelt, A. C., 2016. Adolescent Maturational Transitions in the Prefrontal Cortex and Dopamine Signaling as a Risk Factor for the Development of Obesity and High Fat/High Sugar Diet Induced Cognitive Deficits. Front Behav Neurosci 10, 189.

Reichelt, A. C., Rank, M. M., 2017. The impact of junk foods on the adolescent brain. Birth Defects Res 109, 1649–1658.

Rudebeck, P. H., Walton, M. E., Millette, B. H., Shirley, E., Rushworth, M. F., Bannerman, D. M., 2007. Distinct contributions of frontal areas to emotion and social behaviour in the rat. Eur J Neurosci 26, 2315–2326.

Selimbeyoglu, A., Kim, C. K., Inoue, M., Lee, S. Y., Hong, A. S. O., Kauvar, I., Ramakrishnan, C., Fenno, L. E., Davidson, T. J., Wright, M., Deisseroth, K., 2017. Modulation of prefrontal cortex excitation/inhibition balance rescues social behavior in CNTNAP2-deficient mice. Sci Transl Med 9.

Skelly, M. J., Chappell, A. E., Carter, E., Weiner, J. L., 2015. Adolescent social isolation increases anxiety-like behavior and ethanol intake and impairs fear extinction in adulthood: Possible role of disrupted noradrenergic signaling. Neuropharmacology 97, 149–159.

Spear, L. P., 2000. The adolescent brain and age-related behavioral manifestations. Neurosci Biobehav Rev 24, 417–463.

Sudo, N., Aiba, Y., Oyama, N., Yu, X. N., Matsunaga, M., Koga, Y., Kubo, C., 2004. Dietary nucleic acid and intestinal microbiota synergistically promote a shift in the Th1/Th2 balance toward Th1-skewed immunity. Int Arch Allergy Immunol 135, 132–135.

Takase, K., Tsuneoka, Y., Oda, S., Kuroda, M., Funato, H., 2016. High-fat diet feeding alters olfactory-, social-, and reward-related behaviors of mice independent of obesity. Obesity (Silver Spring) 24, 886–894.

Tanimizu, T., Kenney, J. W., Okano, E., Kadoma, K., Frankland, P. W., Kida, S., 2017. Functional Connectivity of Multiple Brain Regions Required for the Consolidation of Social Recognition Memory. J Neurosci 37, 4103–4116.

Teixeira, D., Cecconello, A. L., Partata, W. A., de Fraga, L. S., Ribeiro, M. F. M., Guedes, R. P., 2017. The metabolic and neuroinflammatory changes induced by consuming a cafeteria diet are age-dependent. Nutr Neurosci, 1–11.

Thaler, J. P., Yi, C. X., Schur, E. A., Guyenet, S. J., Hwang, B. H., Dietrich, M. O., Zhao, X., Sarruf, D. A., Izgur, V., Maravilla, K. R., Nguyen, H. T., Fischer, J. D., Matsen, M. E., Wisse, B. E., Morton, G. J., Horvath, T. L., Baskin, D. G., Tschop, M. H., Schwartz, M. W., 2012. Obesity is associated with hypothalamic injury in rodents and humans. J Clin Invest 122, 153–162.

Trezza, V., Baarendse, P. J., Vanderschuren, L. J., 2010. The pleasures of play: pharmacological insights into social reward mechanisms. Trends Pharmacol Sci 31, 463–469.

Velkoska, E., Warner, F. J., Cole, T. J., Smith, I., Morris, M. J., 2010. Metabolic effects of low dose angiotensin converting enzyme inhibitor in dietary obesity in the rat. Nutr Metab Cardiovasc Dis 20,49–55.

Volkow, N. D., Wang, G. J., Baler, R. D., 2011. Reward, dopamine and the control of food intake: implications for obesity. Trends Cogn Sci 15, 37–46.

Warburton, E. C., Brown, M. W., 2010. Findings from animals concerning when interactions between perirhinal cortex, hippocampus and medial prefrontal cortex are necessary for recognition memory. Neuropsychologia 48, 2262–2272.

Yaseen, A., Shrivastava, K., Zuri, Z., Hatoum, O. A., Maroun, M., 2018. Prefrontal Oxytocin is Involved in Impairments in Prefrontal Plasticity and Social Memory Following Acute Exposure to High Fat Diet in Juvenile Animals. Cereb Cortex.

